# Attentional modulation of persistent visual representations after stimulus offset is coordinated across primary visual and anterior cingulate cortices

**DOI:** 10.64898/2026.10.06.756930

**Authors:** Karen Safaryan, Rohan Bhalodia, Jonathan Shenassa, Peyman Golshani

**Affiliations:** Department of Neurology, David Geffen School of Medicine, University of California, Los Angeles, Los Angeles, CA, USA; Department of Neurobiology, David Geffen School of Medicine, University of California, Los Angeles, Los Angeles, CA, USA; Greater Los Angeles VA Medical Center, Los Angeles, CA, USA; Semel Institute of Neuroscience and Human Behavior, David Geffen School of Medicine, University of California, Los Angeles, Los Angeles, CA, USA; Intellectual and Developmental Disability Research Center, David Geffen School of Medicine, University of California, Los Angeles, Los Angeles, CA, USA; Integrative Center for Learning and Memory, David Geffen School of Medicine, University of California, Los Angeles, Los Angeles, CA, USA

## Abstract

Flexible behavior in complex environments requires animals to infer which sensory inputs are relevant for guiding action and which need to be ignored. Yet, how cognitive control circuits apply contextual knowledge during decision-making is poorly understood. To address this question, we trained head-fixed mice on a cross-modal, context-dependent two-alternative forced-choice (2-AFC) decision-making task. In this task, animals simultaneously received visual and auditory inputs, and the behaviorally relevant modality switched covertly between blocks of trials, requiring animals to infer rule changes from reward feedback alone. Mice successfully adapted their behavior after rule switches and recovered to expert performance. Using simultaneous Neuropixels recordings in the primary visual cortex (V1) and the anterior cingulate cortex (ACC) during task performance, we found that neuronal responses in both regions showed mixed selectivity, multiplexing sensory, choice, and outcome signals. However, V1 neural responses more strongly encoded visual stimulus orientation, while ACC neural responses preferentially encoded outcome. Population decoding further revealed that V1 and ACC neurons maintained a persistent representation of task context throughout the trial, occupying a subspace approximately orthogonal to visual stimulus coding. This contextual representation strengthened progressively as mice adapted their behavior following each switch. Critically, context representations emerged in parallel in V1 and ACC, and decoding accuracies in these two regions were strongly correlated with each other and with behavioral performance across trial blocks. Orthogonal to this shared emergent and sustained context signal, visual stimulus decoding was enhanced when vision was task-relevant. This attentional enhancement in V1 occurred not only during the stimulus presentation period but also after stimulus offset. Similarly, in ACC, a persistent increase of visual stimulus identity encoding in visually relevant session blocks was present after stimulus offset. Taken together, these findings reveal a distributed, inter-areal representation of task context spanning sensory and frontal cortices, suggesting that V1 and ACC contribute to maintaining task-relevant sensory information beyond the immediate sensory response period to provide a circuit-level mechanism for cross-modal attentional control.

## Introduction

To thrive, we must continuously track the outcomes of our decisions and adapt them to environmental changes. Cognitive flexibility requires neural circuits to adapt multimodal information processing as behavioral demands change^1–3^. The connections between sensory and frontal cortical circuits have been studied extensively, but how they contribute to the mechanisms of behavioral flexibility remains elusive.

One well-characterized circuit through which context-dependent decision making may occur is the bidirectional pathway linking the anterior cingulate cortex (ACC) and the primary visual cortex (V1). Anatomical studies have identified projections between visual and prefrontal cortical regions^4,5^, while functional studies indicate that activity in ACC and related prefrontal regions can influence visual cortical processing^6–8^. Recent work further demonstrates that prefrontal feedback to V1 is behavior- and state-dependent, suggesting that top-down signals can selectively gate sensory cortical processing according to the animal’s current behavioral state^8^. Conversely, visual information is transmitted to ACC, where sensory responses can be integrated with task-relevant signals and contribute to goal-directed behavior^9^. Thus, the ACC-V1 circuit provides a potential pathway through which contextual and sensory information can interact bidirectionally. Yet, how contextual and sensory signals are integrated across the two structures and how attentional modulation is implemented before and after the stimulus driving decision-making is poorly understood.

Here, we investigated how contextual information is represented and coordinated across the ACC-V1 circuit during a flexible, cross-modal decision-making task. We trained head-fixed mice on an audiovisual two-alternative forced-choice (2-AFC) task in which the behaviorally relevant sensory modality switched covertly between blocks of trials. Using simultaneous Neuropixels recordings in V1 and ACC, we examined how sensory, choice, and outcome signals are represented within each region, and whether contextual representations emerge with similar timing and dynamics across these two regions. We found that neurons in both areas exhibited mixed selectivity for task variables, but with distinct emphases: V1 responses more strongly encoded stimulus identity, whereas ACC responses more strongly tracked trial outcome. Despite this regional difference in tuning, both areas maintained a persistent representation of task context throughout the trial, with correlated context-decoding dynamics that emerged in parallel as animals adapted their behavior following each rule switch. Furthermore, the behavioral relevance of the visual stimulus enhanced its neural representation beyond the immediate sensory response. These findings indicate that contextual information is represented as a temporally coordinated signal distributed across sensory and frontal cortices; they also point to a circuit-level mechanism by which contextual inference in the associative cortex may extend and sustain task-relevant sensory representations.

## Results

### Mice successfully infer rule switches while performing an audiovisual cross-modal context-dependent decision-making task

We trained head-fixed mice on a cross-modal set-shifting task in which the behaviorally relevant sensory modality alternated covertly between session blocks (Fig. 1A, C). Mice first learned visual and auditory discrimination separately (simple discrimination) before learning a compound discrimination task in which auditory and visual stimuli were presented simultaneously on every trial. Depending on the active decision-making rule (visual context vs. auditory context), either the visual or auditory stimulus determined the correct response, whereas the other modality served as a distractor (Fig. 1A, C).

**Figure 1.**
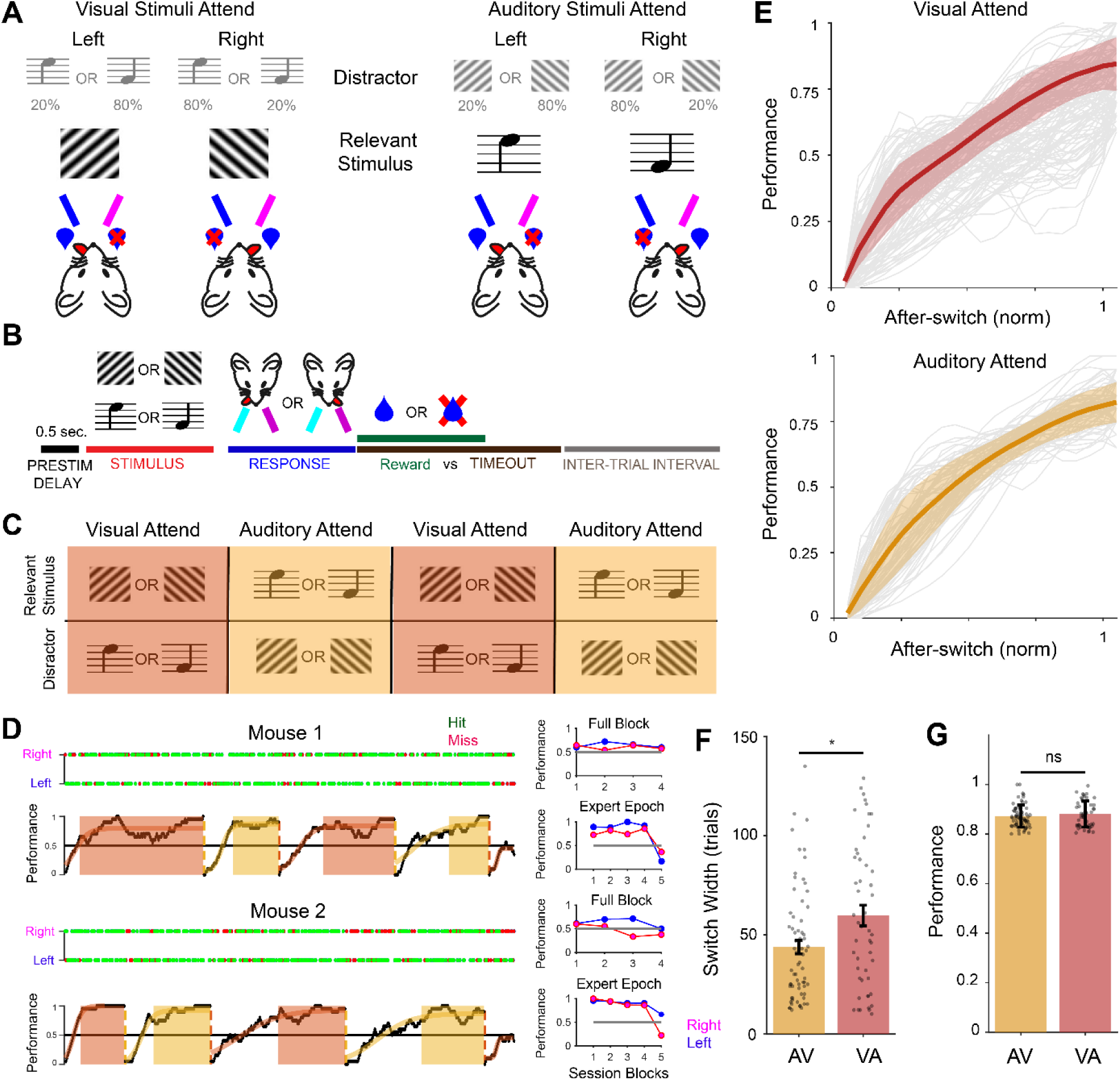
A cross-modal (audio-visual) 2-alternative forced choice (2-AFC) attention-switching task in mice. **(A)** *Task design*. In Visual-Attend blocks (left), mice were rewarded for licking a port that corresponded to the orientation of a visual grating (relevant stimulus) while ignoring a concurrent auditory tone (distractor). In Auditory-Attend blocks (right) the auditory tone determined the rewarded port, and the visual grating was the distractor. In 80% of trials the two stimuli were incongruent (indicating opposite behavioral choices), while 20% of trials combined congruent stimuli. Blue and magenta indicate left/right lick ports; blue droplet/crossed-droplet indicate reward/no-reward outcomes. **(B)** *Trial structure*. A 0.5 sec. pre-stimulus delay is followed by simultaneous presentation of the visual and auditory stimuli (stimulus period: 2 seconds), then a response window (1 second) in which the mouse licked left or right. This is then followed by an outcome period (reward or timeout), and by a variable inter-trial interval. **(C)** *Block structure*. Trials were organized into alternating Visual-Attend (orange) and Auditory-Attend (yellow) blocks; within each block, the relevant and distractor stimulus identities and modalities are indicated. **(D)** Example single-session behavior from two mice. Top: trial-by-trial outcomes (green = hit, red = miss) for right- and left-port choices across the full session. Performance was computed using a sliding window of 10 trials (black) with fitted transition curves (orange/yellow) overlaid on Visual-Attend (orange shading) and Auditory-Attend (yellow shading) blocks, illustrating performance recovery after each switch in stimulus relevance. Right insets: average performance across blocks for the full session (top) and restricted to the expert epoch (bottom) for each mouse with right (magenta) and left (blue) choices shown separately. **(E)** Mean (± SEM, shaded) performance as a function of normalized trial number after a modality switch, for Visual-Attend (left, red) and Auditory-Attend (right, orange) blocks. Thin gray lines show individual session traces. **(F)** Switch width (number of trials needed to reach expert level performance after a block transition) for Auditory-Attend (AV) and Visual-Attend (VA) blocks. Bars show mean ± SEM; dots show individual sessions. Visual-Attend blocks (n = 50) showed significantly wider switches than Auditory-Attend blocks (n=69, *p* < 0.05, rank sum test). **(G)** Expert level performance for AV and VA blocks. Bars show mean ± SEM; dots show individual sessions. No significant difference between conditions (n.s.).

Each trial began with a 500 ms prestimulus delay, where a short sound marked the start of the trial, followed by simultaneous presentation of auditory and visual stimuli for 2 seconds (Fig. 1B). Visual stimuli consisted of drifting gratings at 45° or 135° orientation (spatial frequency: 0.05 cycles per degree; drift speed: 4 Hz; contrast: 100%), and auditory stimuli were 5 kHz or 18 kHz frequency tones. Rule switches occurred after animals’ performance exceeded a threshold of 85% accuracy over 30 trials. Importantly, transitions between auditory-attend (AV) and visual-attend (VA) session blocks were covert and mice had to infer changes in task contingencies from reward feedback and adjust their choices accordingly. This design differs from our previous go/no-go paradigm^10,11^, in which single-modality blocks or brief priming periods preceded each cross-modal switch and gave animals a partial cue to the upcoming context. Here, no such cue was available, so animals had to rely purely on feedback-driven inference.

Expert animals exhibited robust set-shifting behavior across sessions (Fig. 1D). Immediately following covert context switches, performance dropped sharply because animals initially continued to follow the previously relevant sensory modality, which now served as a distractor. Performance gradually recovered after 43±3.4 and 59.5±5.2 subsequent trials for AV and VA session blocks, respectively (Fig. 1F), as mice learned to infer the new stimulus-response contingency. To quantify set-shifting behavior, we aligned performance to block transitions and measured recovery dynamics separately for VA and AV session blocks. In both contexts, performance followed a stereotyped sigmoidal recovery trajectory, reaching expert levels within a comparable number of trials after the switch. Although after-switch dynamics were broadly similar across contexts, VA session blocks required more trials to reach asymptotic performance (Fig. 1E-G). These results indicate that mice can infer task context and flexibly update behavioral strategies according to the currently relevant visual or auditory modality.

### Mixed selectivity with distinct sensory and outcome encoding in simultaneously recorded populations of V1 and ACC

We performed simultaneous Neuropixels recordings in V1 and ACC as mice performed the cross-modal decision-making task. We recorded 541 and 315 units in V1 and ACC, respectively (7 sessions from 5 mice), synchronously with task variables (licks, stimuli, trial onset, context). The units were pooled across all the cortical layers of both structures. These simultaneous V1 and ACC recordings enabled direct, within-animal comparison of context representations across the two regions.

To characterize the distribution of task-variable selectivity across the recorded populations in V1 and ACC (Figure 2A), we fit a generalized linear model (GLM) incorporating both continuous, temporally extended covariates, as well as categorical variables spanning stimulus, choice, and outcome (Figure 2B; see Methods). The model accurately reproduced both the trial-averaged firing rate and the trial-by-trial spiking structure of individual units selective for each of these task variables (Figure 2C).

**Figure 2.**
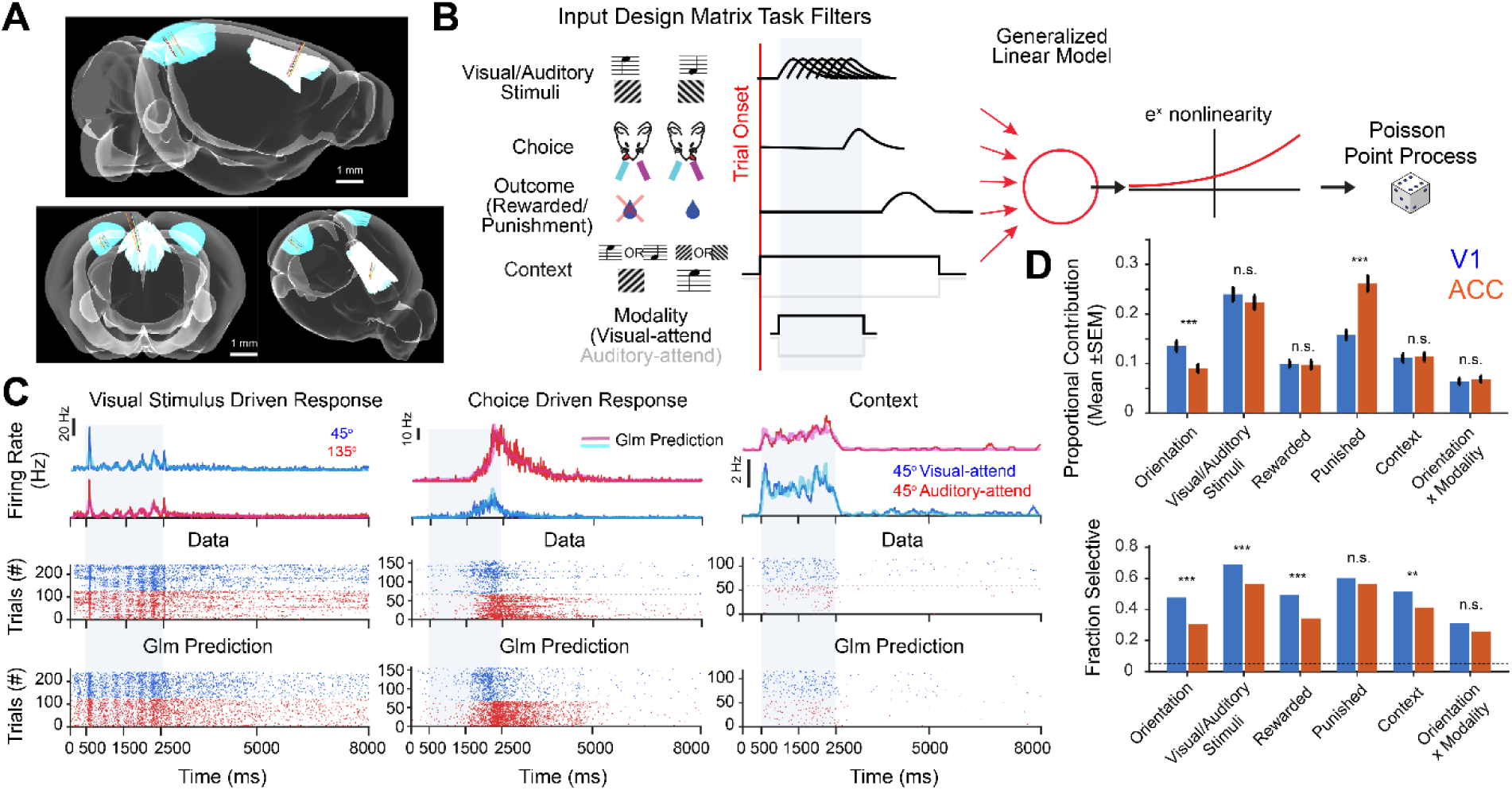
Neuropixels recording from primary visual cortex (V1) and anterior cingulate cortex (ACC) during task performance demonstrates mixed selectivity of responses in both structures. **(A)** Reconstruction of Neuropixels probe tracks, registered to the Allen Mouse Brain Common Coordinate Framework (10 µm resolution), shown in sagittal and coronal views highlighting probe placement in visual cortex (blue) and anterior cingulate cortex (green). **(B)** Overview of the generalized linear model (GLM). Spike trains were aligned to trial events and convolved with a set of event-locked kernels (basis functions) representing task covariates (stimulus, choice, and outcome-related variables), producing a design matrix that was combined linearly and passed through an exponential nonlinearity to generate the model’s instantaneous predicted firing rate (Poisson GLM). **(C)** Validation of model predictions for three representative example units, driven respectively by stimulus identity (45° vs. 135°, left), choice (middle), and context (45° visual-attend vs. 45° auditory-attend, right). For each: top, trial-averaged PSTH split by group, with the GLM-predicted rate overlaid as a dashed line; middle, raw single-trial spike raster for the same trials, stacked by group; bottom, spike trains sampled from the GLM’s predicted rate for the corresponding trials. **(D)** GLM results across the recorded population in V1 (blue, n = 541 units) and ACC (orange, n = 315 units), for each task covariate: proportional contribution (mean ± SEM) to the model’s predicted firing rate (top), and the fraction of units with a statistically significant contribution relative to a chance-level threshold (dashed line, bottom). Asterisks indicate a significant V1-vs-ACC difference for that covariate (Welch’s t-test on proportional contribution; two-proportion z-test on fraction selective; *** p < 0.001, ** p < 0.01, * p < 0.05, n.s. = not significant).

Across the population, V1 and ACC showed distinct profiles of task-variable encoding (Fig. 2D). Consistent with its role in early sensory processing, activity in V1 units showed a significantly larger proportional contribution from visual stimulus orientation than ACC units (Fig. 2D, p < 0.001). In addition, a significantly greater fraction of units in V1 were selective for visual stimulus orientation compared to ACC (Fig. 2D, p < 0.001). In contrast, ACC units showed a significantly larger proportional contribution from punishment outcome than V1 (p < 0.001), despite showing a similar fraction of significantly punishment-selective units. This indicated that, on average, punishment signals contributed more strongly to the modeled firing rate of individual ACC units. The proportional contribution of visual/auditory stimulus identity, rewarded outcome, and the orientation × modality interaction did not differ significantly between regions. These results suggest that while neurons in both regions multiplex sensory, choice, and outcome signals, V1 more robustly encodes stimulus-related variables at the single-unit level, whereas ACC neurons more selectively respond to the outcome (specifically the punishment) of the animal.

### Visual stimulus identity encoding in V1 is enhanced during and after the stimulus presentation period in visual-attend blocks

To determine whether behavioral context influences sensory representations in V1, we examined the encoding of visual stimulus identity during the performance of the audio-visual cross-modal decision-making task. We trained linear decoders to classify visual stimulus identity from population activity throughout the AV and VA trial blocks (Fig. 3A–D).

**Figure 3.**
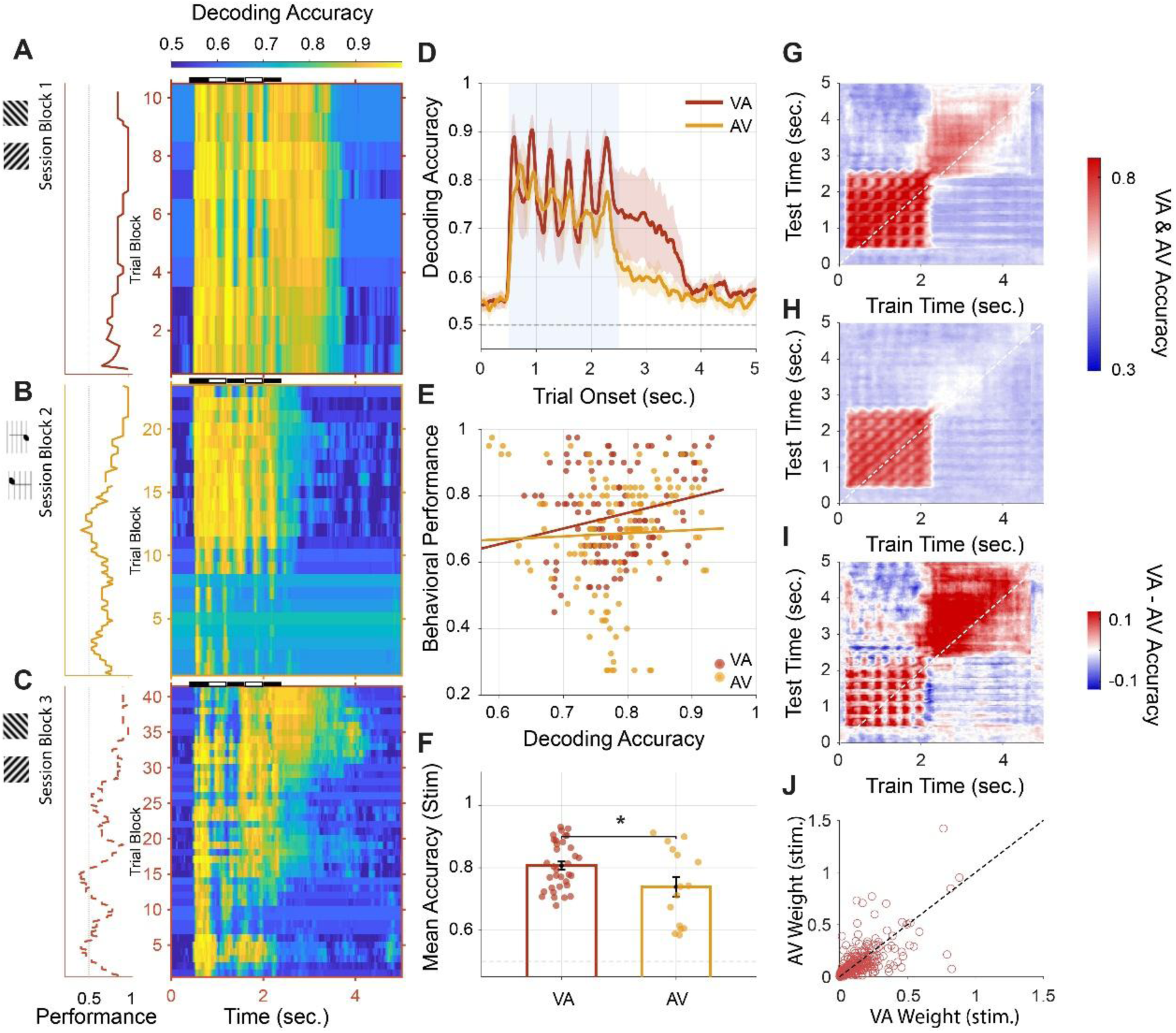
Task context enhances the strength and stability of visual representations in V1. **(A–C)** Single-session decoding heatmaps. Each panel shows trial-by-trial decoding accuracy of the visual stimulus across time within a single example session, for three successive task blocks: Block 1 (VA context, visual-relevant; dark red), Block 2 (AV context, auditory-relevant, orange), and Block 3 (VA context, dashed dark red). Color indicates decoding accuracy. Performance strips to the left of each heatmap show the animal’s behavioral hit rate across batches for the corresponding block. **(D)** Grand-mean decoding accuracy as a function of time within the trial, averaged across all expert-phase batches (hit rate ≥ 75%) and all sessions (n = 7), separately for VA (dark red, solid) and AV (orange, solid) first blocks and VA2 (dark red, dashed) and AV2 (dark olive, dash-dot) second blocks where available. Shading indicates ± SEM across sessions. The blue-shaded region (0.5–2.5 s) marks the early stimulus period. Dashed horizontal line indicates chance (0.5). **(E)** Scatter of behavioral performance against neural decoding accuracy across batches separately for VA (dark red, r = 0.2317, p = 0.008) and AV (orange, r = 0.0401, p = 0.6366) across all sessions. Each dot is one batch from one session. **(F)** Mean decoding accuracy during the stimulus presentation period (0.5–2.5 s) for VA and AV contexts, pooled across blocks. Open bars show group means ± SEM. Individual session values are shown as filled dots (* p = 0.0110, one-tailed t-test). **(G-I)** Cross-temporal generalization matrices for visual stimulus decoding. Each matrix shows the accuracy of a classifier trained at one time point (x-axis, train time) and tested at every other time point (y-axis, test time), averaged across expert-phase batches and sessions. (G) VA context: broad off-diagonal structure indicates a temporally stable population code for visual stimulus identity during visual-relevant behavior. (H) AV context: reduced off-diagonal structure reflects weaker and less sustained visual representations when vision is behaviorally irrelevant. (I) Difference matrix (VA − AV): positive values (red) indicate time-point pairs where visual representations generalize more strongly in the VA context, revealing both stronger initial encoding and greater temporal persistence when vision guides behavior. **(J)** Visual stimulus decoding weights for VA and AV session blocks across units (r = 0.73, p = 0). Weights are systematically larger in VA than AV context (p = 0.043).

To track how the neural representation of the visual stimulus changed as mice adapted within each modality-relevant block, we used a moving-window decoding approach. Trials were grouped into overlapping windows of 40 consecutive trials and each new window shifted forward by 5 trials so that we could follow decoding accuracy smoothly as behavior evolved across VA and AV session blocks. For each window, we trained a linear classifier (support vector machine, SVM) to distinguish the two grating orientations (45° vs. 135°) from the simultaneous activity of all recorded V1 units, spanning all cortical layers and including both putative excitatory and inhibitory neurons. This classifier was applied independently at every 10 ms time bin spanning the full trial time, giving a moment-by-moment readout of how well stimulus identity could be decoded at each point in the trial, for each trial block. The classifier was always tested on trials it had not been trained on. Repeating this across successive trial blocks produced a two-dimensional map of decoding accuracy for each session block (Figs. 3A–C): one axis tracking progress across trials within a session block, the other tracking time within a trial. To assess the overall time course during periods of stable expert performance, we averaged decoding accuracies across all expert-phase trial blocks (performance ≥85%) separately for VA and AV session blocks (Fig. 3D). Decoder weights from each VA and AV block were also retained and averaged across expert-phase trial blocks to characterize the neural subspace supporting visual stimulus coding in each context (Fig. 3J). Within individual sessions, decoding accuracy was higher during periods of expert performance in VA session blocks (Fig. 3A)–where mice based their decisions on visual stimuli–compared to in AV session blocks. Following covert switches into AV session blocks, decoding accuracy dropped abruptly, despite unchanged sensory input (Fig. 3B), before gradually recovering as animals adapted to the new task demands. A subsequent switch back to the VA context produced a similar pattern (Fig. 3C).

Consistent with this observation, decoding accuracy closely followed behavioral performance across sessions and contexts (Fig. 3E). Trial blocks associated with higher task performance exhibited stronger visual stimulus decoding, revealing a strong relationship between sensory representations in V1 and the animal’s behavior. Notably, when analysis was restricted to epochs of expert performance, visual stimulus decoding remained significantly stronger during VA session blocks than during AV session blocks (Fig. 3D, F), despite identical visual stimuli. These results indicate that the behavioral relevance of visual information enhances its neural representation within V1. Moreover, the visual stimulus identity can be reliably decoded during the subsequent choice-related (stimulus-offset) period only in VA session blocks (Fig. 3D).

We next asked whether the behavioral relevance of the visual stimuli also influenced the temporal organization of their neural representations during and after the stimulus periods. To assess the temporal stability of visual stimulus representations, we applied cross-temporal generalization decoding, which asks whether visual representations stay the same or change from moment to moment. This produces a time-by-time map in which the diagonal shows the results of the decoder trained and tested at the same timepoint and the off-diagonal shows how well a decoder trained at one timepoint can decode visual stimuli at other timepoints. Cross-temporal decoding analyses revealed sustained generalization across the stimulus period during both VA and AV session blocks, consistent with a stable population code for visual stimulus identity (Fig. 3G, H). In VA session blocks, this generalization extended into the stimulus-offset period (Fig. 3G), consistent with the persistent decoding accuracy described above. This persistent stimulus-offset encoding was absent in AV session blocks (Fig. 3H), where decoding accuracy declined once the stimulus disappeared. Direct comparison of the two contexts confirmed stronger and more persistent decoding in VA blocks throughout the stimulus and choice task periods (Fig. 3I).

Despite these differences in decoding performance between VA and AV session blocks, the neuronal populations contributing to visual stimulus decoding were largely shared across both conditions during the stimulus period. Decoder weights were strongly correlated between VA and AV blocks (Fig. 3J), indicating that similar neurons participated in encoding visual information regardless of which modality was task-relevant. Notably, decoding accuracy remained high beyond stimulus offset specifically in VA blocks. Together, these findings demonstrate that visual representations in V1 are dynamically shaped by behavioral relevance, becoming stronger and more temporally stable when visual information is relevant for guiding behavior.

### Primary visual cortex encodes task context as a sustained population representation

To determine whether V1 populations represent the relevant task context, we trained linear decoders to distinguish VA and AV session blocks using neural activity recorded during periods of expert performance with an equal number of trials from VA and AV session blocks. All other aspects of the approach—moving windows, time-resolved training and testing, and cross-validation—matched the stimulus-decoding analysis described above.

Within individual session blocks, context decoding accuracy increased progressively across successive trial blocks following a rule change (Fig. 4A, B). Immediately after a covert switch, contextual information was relatively weak, but decoder performance steadily improved as animals adapted to the newly relevant modality. This gradual increase was observed following transitions into both VA and AV blocks (Fig. 4C). Overall, these results indicate that V1 contextual representations are dynamically updated in parallel with behavioral adaptation.

**Figure 4.**
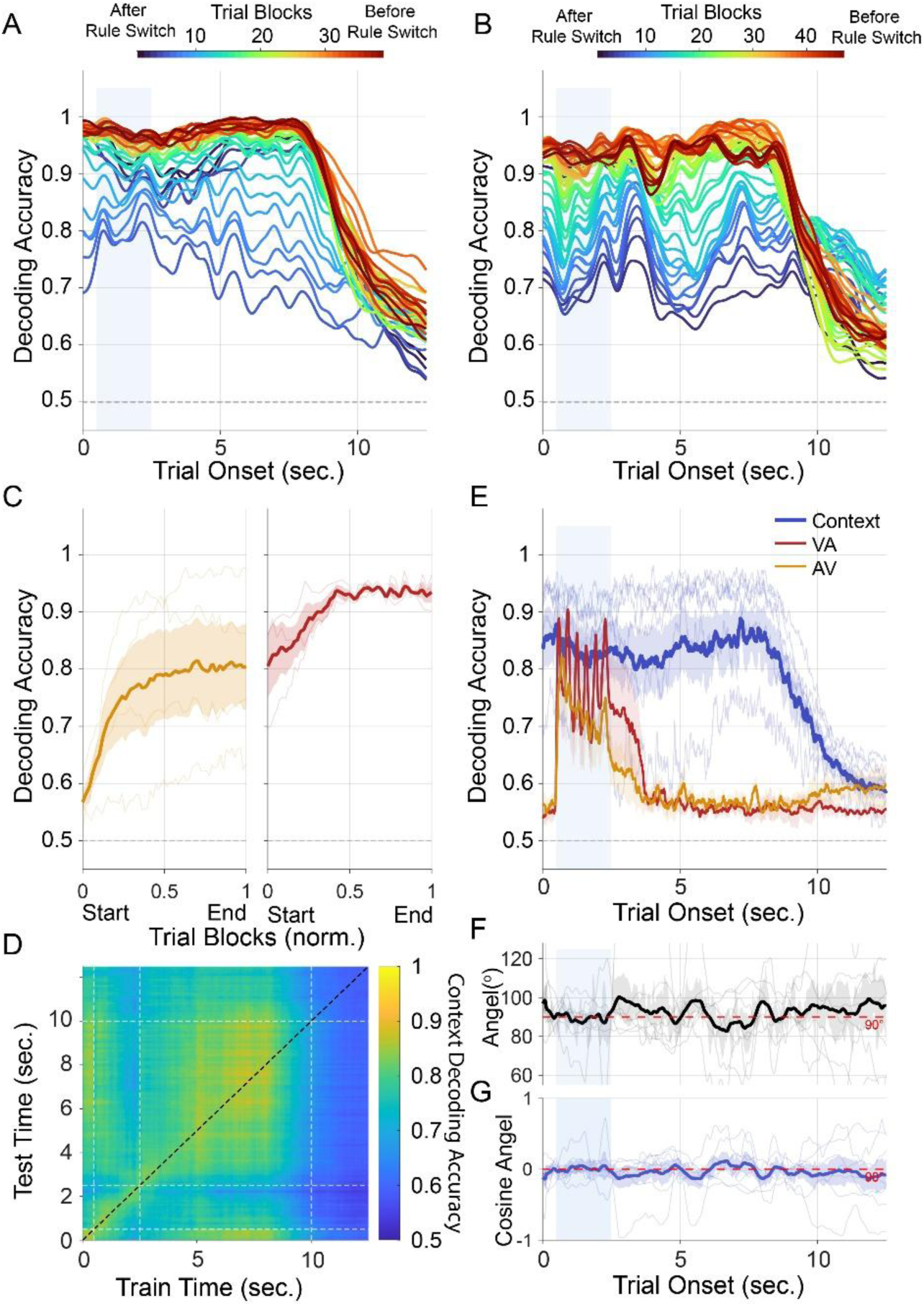
V1 simultaneously encodes task context and visual stimulus identity in dynamically coupled, approximately orthogonal subspaces. **(A–B)** Within block emergence of contextual representations. Each line shows the time-resolved accuracy of a context decoder applied to successive batches within a single example session, for the visual-attend (A) and auditory-attend (B) blocks. Line color indicates batch number within the block, progressing from early (blue/cool) to late (red/warm) batches. **(C)** Contextual decoding accuracy improves across trials from start to end of both visual (right) and auditory (left) task blocks. Thin lines present individual sessions, shaded regions denote ±1 SEM. **(D)** Cross-temporal decoding generalization showing contextual decoding accuracy, where a classifier trained at each time point is tested at every other time point. **(E)** Simultaneous multiplexed encoding of context and visual stimulus identity. Grand-mean decoding accuracy (n=7 sessions) for three simultaneously trained decoders: context (VA vs AV, blue), visual stimulus identity during VA blocks (red), and visual stimulus identity during AV blocks (orange). Thin lines show individual session traces, shaded regions denote ±1 SEM. **(F-G)** Visual–Context hyperplane angle (F) and similarity (G) across overall trial time. Thin lines show individual sessions; shaded region denotes ±1 SEM.

We next examined the temporal organization of contextual representations using the same time-shifted decoding approach used in Figure 3G-I, now applied to context rather than to the visual stimulus. A decoder was built at one reference point in the trial and then tested at every other time point, using held-out trials pooled across the session’s VA and AV blocks. While contextual representation in the V1 was relatively stable during the post-stimulus task period, the decoder revealed an overall weaker stability and generalization of contextual representation compared to visual stimulus (Fig. 4D). Unlike visual stimulus representations, which were largely restricted to the stimulus presentation period (Fig. 3G–I), contextual information was present throughout the trial. Consistent with this, context decoding rose before stimulus onset, remained elevated throughout stimulus presentation, and persisted into the post-stimulus period (Fig. 4E).

Despite overlapping in time, the context and visual signals occupied largely separate directions within the population activity space. By separating hyperplanes (see Methods) for the visual stimulus and context decoders, we found that the angle between these decoding directions stayed close to 90° throughout the trial (Fig. 4F) and their cosine similarity remained near zero (Fig. 4G), confirming that context and sensory information were encoded along largely independent (orthogonal) dimensions.

These findings demonstrate that V1 maintains a persistent representation of task context that evolves with behavioral adaptation and coexists with sensory encoding in an orthogonal population subspace. Such an organization may allow V1 to simultaneously represent incoming sensory information and the behavioral framework within which that information must be interpreted.

### Context representations emerge in parallel across V1 and ACC during behavioral adaptation

The presence of robust contextual signals in V1 raised the question of whether these representations are coordinated with higher-order cortical areas implicated in cognitive control and behavioral flexibility. To address this question, we examined task-context representations in the ACC, a region implicated in rule-guided behavior.

Using the same decoding framework applied to V1, we trained linear classifiers to distinguish VA and AV contexts from ACC population activity recorded during periods of expert performance. To facilitate direct comparisons between regions, decoding analyses were performed using matched population sizes in ACC and V1. We then applied the decoders independently to each trial block within each session block in order to examine the evolution of contextual representations following covert rule switches.

Similarly to V1, contextual representation accuracy in ACC increased progressively across successive trial blocks following a rule switch (Fig. 5A, B). Immediately after a covert switch, context decoding accuracy was relatively weak in both regions but gradually increased as mice started attending to the newly relevant sensory modality. This pattern was observed following transitions into both AV and VA session blocks, indicating that contextual representations in ACC and V1 evolve dynamically during behavioral adaptation.

**Figure 5.**
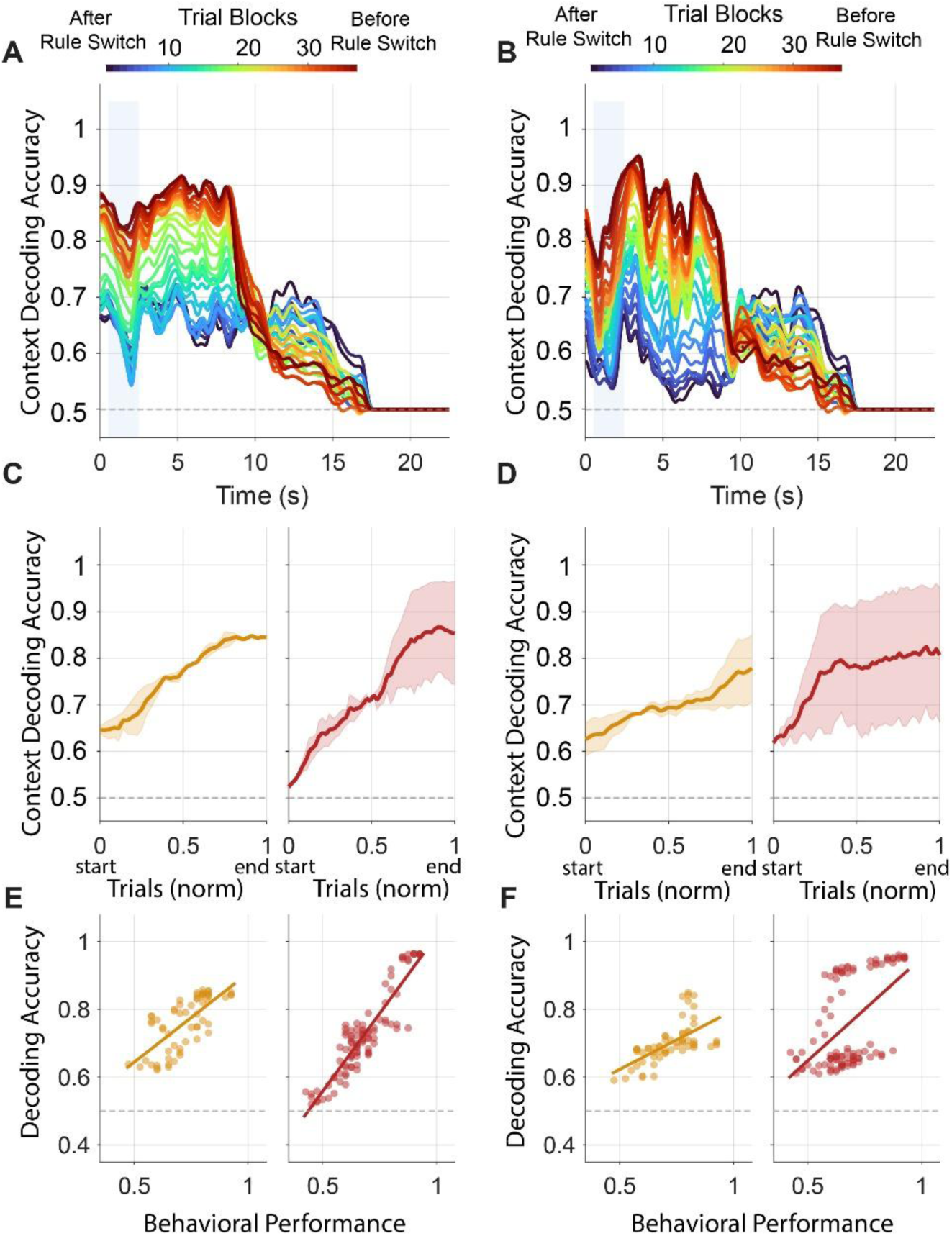
Context representations emerge in parallel across V1 and ACC during behavioral adaptation. **(A–B)** Within block emergence of contextual representations in V1 (A) and ACC (B). Each trace shows trial time-resolved context decoding accuracy for a successive batch of trials from a representative session. Color indicates batch number within the block, progressing from early (blue) to late (red) stages of adaptation following a context switch. **(C-D)** Contextual decoding accuracy improves across trials from start to end of both visual (right) and auditory (left) task blocks for V1 and ACC. Decoding accuracy is plotted as a function of normalized trial position within each block. Thin lines present individual sessions, shaded regions denote ±1 SEM. **(E-F)** Relationship between behavioral performance and context decoding accuracy in V1 (E) and ACC (F). Each point represents a trial batch from one session. Lines are linear regression fits.

To quantify these dynamics across sessions, we aligned decoding accuracy to session block transitions and examined this evolution. Context decoding increased steadily from the beginning to the end of both AV and VA session blocks in V1 and ACC (Fig. 5C, D). Thus, contextual information accumulated progressively in both sensory and frontal cortical populations as animals inferred the currently relevant task rule.

Because contextual representations emerged in parallel with behavioral recovery, we next examined the relationship between decoder performance and behavioral performance. In both V1 and ACC, context decoding accuracy was strongly correlated with behavioral accuracy across trial blocks (Fig. 5E, F), indicating that neural representations of task context closely tracked the animal’s behavioral state. These findings suggest that contextual signals in both cortical regions reflect the ongoing process of rule acquisition and maintenance.

Hence, contextual representations are not confined to V1, but rather emerge in parallel across frontal and sensory cortical populations, forming a distributed network that tracks behavioral adaptation during flexible decision making.

### Differential contextual modulation of post-stimulus period activity

Visual stimulus identity was decoded from population activity during VA and AV session blocks separately (see Methods). This allowed us to isolate the effects of variable attention levels between VA and AV session blocks from sensory representation.

During the visually-evoked response (0.5 – 2.5 s following trial onset), decoding accuracy was similarly high between VA and AV session blocks in both V1 and ACC (Fig. 6A–C), indicating that the initial sensory-driven representation of the stimulus was comparable regardless of its behavioral relevance. Following stimulus offset, however, the two regions diverged in how this representation evolved. In V1, the elevated VA decoding accuracy declined gradually after stimulus offset, approaching AV levels by approximately 4 s after trial onset. In ACC, by contrast, the VA−AV difference persisted at a reduced but comparatively stable level up to 6 s after trial onset (Fig. 6C). Averaging decoding accuracy within an early post-stimulus window (2.5–4 s, chosen to capture the period of peak divergence in both regions) confirmed this pattern: attentional modulation of visual-stimulus decoding accuracy was statistically significant in both V1 (unpaired t-test, p = 5.4*10^-11^ Bonferroni-corrected; Wilcoxon rank-sum p = 3.41*10^-8^, for trial blocks with ≥85% performance) and ACC (unpaired t-test, p =0.0096, corrected p = 0.019, rank-sum 0.014) with higher decoding accuracy during VA than AV session blocks in both regions (Fig. 6D).

**Figure 6.**
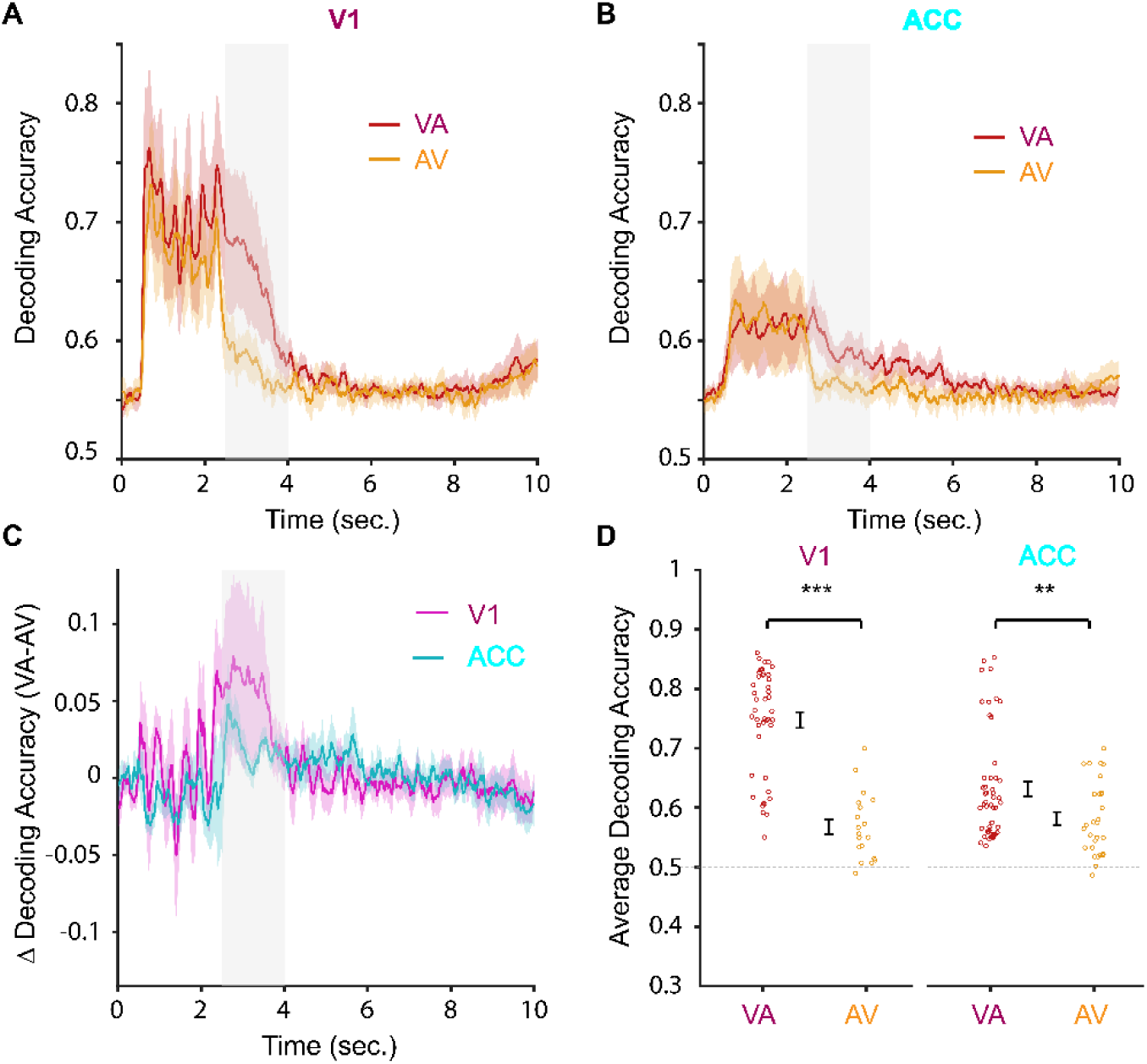
Attentional modulation of visual stimulus decoding is transient in V1 and sustained in ACC. **(A–B)** Population decoding accuracy for visual stimulus identity (left vs. right grating), shown separately for trials in which the visual stimulus was task-relevant (Visual-Attend, VA; red) versus task-irrelevant (Auditory-Attend, AV; gold), in (A) V1 (n = 44 for VA and n = 19 for AV trial blocks) and (B) ACC (n = 51 for VA and n = 29 for AV trial blocks). Stimulus identity was defined by the physical grating orientation regardless of behavioral relevance, so VA and AV trials differ only in attentional state, not stimulus modality. Traces show mean ± SEM decoding accuracy across sessions. **(C)** Attentional modulation of visual decoding (VA − AV difference), for V1 (magenta) and ACC (cyan), computed by first averaging VA and AV time courses within each session and then differencing (preserving the natural session-level pairing), followed by averaging across sessions (mean ± SEM). Both areas show comparable modulation during the visually-evoked response (0–2.5 s). After stimulus offset, V1’s modulation declines toward baseline within ∼1–1.5 s, while ACC’s shows a smaller, more temporally extended positive deflection, descriptively peaking around 3.3–3.8 s post-stimulus. **(D)** Average decoding accuracy in a 2.5–4 s post-stimulus window, per session, for VA and AV trials (error bars, mean ± SEM). This window-averaged comparison revealed significant modulation in both V1 (unpaired t-test, p = 5.4*10^-11^, Bonferroni-corrected) and ACC (unpaired t-test, p =0.019, Bonferroni-corrected). *** indicate significance level 0.01 or less.

These results suggest that while V1 and ACC initially represent the visual stimulus with comparable, context-independent fidelity, the two regions differ in how that representation is maintained after the stimulus disappears. Strong contextual modulation of visual stimulus is present in V1 and ACC during the post-stimulus period. V1’s contextual enhancement is transient, whereas ACC’s is smaller in magnitude but more temporally sustained—suggesting a role for ACC in maintaining task-relevant sensory information beyond the period of direct stimulus drive.

## Discussion

Here, we used simultaneous Neuropixels recordings in V1 and ACC to examine how contextual and sensory information are represented and coordinated within a distributed sensory–frontal circuit during a flexible, cross-modal decision-making task. Although neurons in both regions exhibited mixed selectivity for stimulus, choice, and outcome variables, V1 more strongly and more frequently encoded stimulus orientation, whereas ACC units showed a disproportionately larger contribution of punishment signal. Both V1 and ACC maintained a persistent representation of task context that emerged progressively following each covert rule switch. The decoding accuracies in the two regions were correlated with each other and with behavioral performance. Remarkably, the behavioral relevance of the visual stimulus enhanced its neural representation not only during the sensory response itself but also after stimulus offset. Collectively, these findings support a network model in which contextual inference is implemented as a coordinated, inter-areal computation spanning sensory and frontal cortices.

A central finding of this study is that context representations emerged with similar dynamics and were correlated in strength across V1 and ACC as animals adapted to each covert rule switch. This further extends our previous findings^10^ on contextual representation in V1 occurring in a broader, distributed network framework. This is consistent with a growing body of evidence for a distributed model in which contextual information is shared^12,13^ between sensory and frontal populations, potentially through the wider reciprocal ACC-V1 projections described anatomically and functionally in prior work^6–9^. The tight correlation between context-decoding accuracy and ongoing behavioral performance in both regions further suggests that these signals track—and may contribute to—the animal’s evolving internal model of the task.

Within V1, the context signal occupied a subspace that remained close to orthogonal to the subspace supporting visual stimulus decoding throughout the trial. This organization is consistent with our previous work^10^ and other reports showing that orthogonal coding geometries allow neural populations to represent multiple task variables simultaneously while minimizing mutual interference. Different task sets^14^, however, often rely on overlapping rather than fully segregated neural representations. Shared neural subspaces can provide an efficient substrate for reusing neural computations across tasks^15,16^, but the same overlap can also create interference when competing task demands must be maintained or rapidly switched^14,17,18^. In set-shifting tasks, a new task set can transiently enhance representation of both relevant and irrelevant sensory cues^14,17^. However, with increasing task certainty, representations of task-relevant stimuli become progressively more distinguishable at the population level^19–21^, while representations of concurrently irrelevant cues can be attenuated over the same time course ^22–25^. Importantly, these processes often need not be confined to a single cortical region. Sensory cue categorization, task identity, and choice information are distributed across multiple cortical areas, with representations that are partially shared across the cortical hierarchy rather than strictly segregated by function^26–30^.

The most consistent signature of this population-level encoding in our results is the attentional enhancement of behaviorally relevant sensory cues. However, the most notable result of this study is the persistence of attentional enhancement of visual stimulus decoding following stimulus offset. Both V1 and ACC showed comparable attentional modulation during the visually-evoked response, indicating that the initial gain in sensory representation associated with task relevance is shared across the two areas. After the stimulus disappeared, this enhancement was temporally maintained in V1 and ACC to varying degrees. V1 displayed a gradual decline in stimulus representation following stimulus offset, while ACC, or the coordinated V1-ACC circuit, sustained task-relevant sensory information well after direct sensory drive ended.

Given known top-down projections from ACC and related prefrontal regions back to V1, and evidence that this feedback is state- and behavior-dependent^3,8^, one interpretation is that ACC’s persistent representation of the recently presented, behaviorally relevant stimulus reflects a working-memory-like maintenance signal that could, in principle, be relayed back to V1 to prolong sensory-related activity there. An alternative possibility is that this maintained representation is generated largely within ACC itself based on input received during the stimulus period without requiring continued engagement of V1. Our correlational decoding approach cannot distinguish between a feedback-dependent extension of the V1 representation and an independently maintained ACC trace. Future studies using causal circuit manipulations—for example, optogenetic suppression of ACC-to-V1 feedback restricted to the post-stimulus window—will be needed to test whether the sustained ACC signal is necessary for extending sensory-related processing in V1, or whether it instead reflects a parallel, ACC-intrinsic memory trace.

More broadly, models of contextual computation typically implicate mechanisms distributed across multiple brain regions^12,13^. Whether, and to what extent, early sensory cortical regions participate in this computation remains debated^19,28,31^. Engel et al. (2015) offered a theoretical account of how downstream, choice-related signals could nonetheless come to reshape sensory tuning over learning^32^. Stable category representations emerge because they carry choice-correlated activity, arising from the plasticity of top-down projections originating in decision-related neurons. Experimentally, mouse visual cortical areas have been shown to develop category tuning with learning at early stages of the visual processing hierarchy, indicating that abstract, semantic content can be represented outside canonical association cortices^31^.

Our data offers a candidate substrate for such a mechanism operating on V1 itself. Visual stimulus identity remained decodable and attentionally modulated in V1 through the choice period, only gradually returning to baseline several seconds after stimulus offset (Fig. 6). In ACC, this same attentional modulation was smaller in magnitude but significantly more persistent across the same post-offset window (Fig. 6A, B). This extended, choice/outcome-adjacent window of coordinated V1-ACC engagement—layered onto the orthogonal but temporally parallel context representations we observe in both regions throughout the trial—provides a plausible window during which ACC output can modulate V1. Repeated across trials, such feedback could progressively retune V1 responses toward the animal’s learned category structure^33,34^. This may explain how contextual information can be represented in early sensory cortices without originating from these structures.

## Materials and Methods

### Animals and surgery

All experimental procedures were approved by the University of California, Los Angeles Office for Animal Research Oversight and by the Chancellor’s Animal Research Committees. We used 7–10-week-old male and female B6129SF1/J mice (F1 hybrids from a cross between C57BL/6J females and 129S1/SvImJ males; The Jackson Laboratory). Mice were injected with a systemic analgesic (carprofen, 5mg/kg body weight, intraperitoneal) and after 10 minutes were anesthetized with isoflurane (3–5% induction, 1.5% maintenance). They were then placed in a stereotaxic frame. Body temperature was maintained at 37°C throughout using a feedback-controlled heating pad (Harvard Apparatus). Pressure points and incision sites were injected with lidocaine (2%), and eyes were protected from desiccation using artificial tear ointment. The surgical site was sterilized with iodine and ethanol, the scalp was incised and removed, and a custom-made, lightweight, omega-shaped stainless-steel head holder was implanted on the skull using Vetbond (3M) and dental cement (Ortho-Jet, Lang). A recording chamber was built from dental cement around the future craniotomy sites and the skull was covered and sealed with a silicone elastomer sealant (Kwik-Sil, WPI). Mice recovered from surgery and received carprofen (5 mg/kg body weight, intraperitoneal injection) for 2 days daily and amoxicillin (0.25 mg/mL in drinking water) for 7 days post-operatively. Following recovery, mice were water-restricted to 90 percent of their body weight and trained on the behavioral task described below.

### Electrophysiology surgery

Approximately 24 h before electrophysiological recordings, mice were anesthetized with isoflurane and a small craniotomy (0.5 mm diameter) was made above the right cerebellum; a silver chloride ground wire was implanted in the craniotomy and fixed in place with dental cement. Circular craniotomies (1 mm diameter) were made above ACC (1.8 mm anterior to bregma, 0.25 mm right of the midline) and V1 (4.2 mm posterior to bregma, 2.5 mm lateral to midline, right hemisphere). The exposed skull and brain were covered and sealed with a silicone elastomer sealant (Kwik-Sil, WPI). On the day of recording, the mouse was placed in the recording apparatus and the head bar was fixed to a post.

### Behavioral task and training

#### Apparatus

Water-restricted, head-fixed mice were trained and tested in a custom behavioral rig equipped with a 9.7-inch retina display (2048 × 1536 pixels) placed 25 cm eye-to-screen distance with a viewing angle of 45° from the center of the monitor to the left eye of the mouse for presentation of drifting visual grating stimuli. Visual stimuli were generated and controlled using the Psychophysics Toolbox^35^ in Matlab. Auditory tone were delivered from both left and right speakers at 70 dB. The servo motors moved two lick spouts into close proximity of the animal’s mouth 250 ms before stimulus presentation ends. Mice were rewarded after the second lick on the same side spout. The other spout was moved out of reach after choice was made^36^. The rig was enclosed in a soundproofed box to mitigate environmental interference. Licks were detected and recorded using either capacitive or optical lick-detection sensors. Correct responses were rewarded with 3 μL of water, whereas incorrect responses were followed by a 3-s timeout before the next trial.

#### Task design

Mice performed a cross-modal, context-dependent two-alternative forced-choice (2-AFC) task in which the behaviorally relevant sensory modality (vision or auditory) switched covertly across blocks of trials (Fig. 1A, C). On each trial, mice were presented with both a visual grating stimulus (45° or 135° orientation; drifting at a temporal frequency of 4 Hz, spatial frequency of 0.05 cycles/degree, 100% contrast, 2 seconds in duration) and an auditory tone (5 kHz or 18 kHz, 70dB, 2 seconds in duration), with each stimulus independently indicating that the mouse should lick the right or the left spout for water. During Visual-Attend (VA) blocks, the correct response was determined by the orientation of the visual grating while the auditory tone served as a task-irrelevant distractor; during Auditory-Attend (AV) blocks, tone identity determined the correct response and the visual grating served as the distractor (Fig. 1A). On 80% of trials, the two stimuli were incongruent (each independently indicating opposite responses), while on the remaining 20% of trials, the two stimuli were congruent (both indicating the same response).

Each trial began with a 500 ms pre-stimulus delay, during which a brief tone marked trial onset, followed by simultaneous presentation of the visual and auditory stimuli for 2 s (Fig. 1B). A 1 s response window followed stimulus offset, during which the animal reported its choice with a left or right lick. Correct choices were rewarded with a water droplet from the same spout; incorrect choices resulted in a punishment timeout period lasting 3 seconds. Random Inter-Trial Intervals (ITIs) were derived from a gamma distribution.

The behaviorally relevant modality switched between VA and AV blocks once the animal’s performance reached or exceeded 85% accuracy over a moving window of 30 trials. Modality switches were entirely covert. No external cues signaled the transition, so mice had to infer the rule change purely from the reward feedback on subsequent trials and adjust their choices accordingly. Expert mice typically made 3-4 switches per session.

#### Training protocol

Mice were trained in three sequential phases. In the first phase (simple discrimination), mice learned to discriminate the visual and auditory stimuli separately, in dedicated unimodal sessions, until reaching 85–90% accuracy on 2–3 consecutive days. In the second phase (compound discrimination), auditory and visual stimuli were presented simultaneously on every trial using 80% incongruent and 20% congruent trials. In the third and final phase (context-switching), the 80% incongruent / 20% congruent trial structure remained in place while the covert modality switches described above were introduced. Mice were trained until they exhibited reliable set-shifting behavior—recovering to expert performance after each covert switch (see Results)— before undergoing electrophysiology surgery.

### Neuropixels recording

Recordings were performed in awake, head-fixed mice. The head-fixed animals were placed on the rig and the silicone sealants covering the craniotomies were removed. A 0.9% NaCl solution was applied throughout the recording to keep the brain surface wet. A Neuropixels 1.0 probe (IMEC)^37^ was referenced to the brain surface and probes were mounted on a New Scale Pathway MPM micromanipulator system, operated through Pathway MPM Control software, and lowered slowly (∼1 min/mm) to a final depth of 3.0 mm, spanning all cortical layers of V1 and oriented orthogonal to the columnar axis of ACC. Probes were dipped in CM-DiI, DiD, or DiO (Thermo Fisher Scientific, cat. no. V22889) before each insertion to allow post hoc histological reconstruction of probe tracks. In each session, up to two probes were inserted simultaneously (one in V1, one in ACC), each at a fixed angle within the coronal plane 10° and −10° from vertical for V1 and ACC, respectively. We recorded up to five sessions per mouse.

Electrophysiological signals and trial-start timestamps were acquired at 30 kHz using SpikeGLX with an IMEC Neuropixels PXIe module installed in a National Instruments PXIe-1071 chassis (LFP was acquired at 2.5 kHz sampling frequency).

### Histology, probe tracking, and imaging

At the end of the recording period, mice were deeply anesthetized with isoflurane and transcardially perfused with 0.1 M Phosphate Buffer Solution (PBS) followed by 4% paraformaldehyde (PFA). Brains were dissected, post-fixed in 4% PFA overnight at 4°C, and cut into 100-µm coronal sections on a vibratome (VT1200S, Leica Biosystems). Sections were mounted on SuperFrost Plus slides (VWR) with VECTASHIELD mounting medium (Vector Laboratories). Because probes were inserted at fixed angles within the coronal plane, the CM-DiI-, DiD-, and DiO-labeled tracks could be followed across consecutive coronal sections. Sections were imaged on a confocal microscope (ApoTome2 microscope, Zen2 software, Zeiss) with a ×10 objective, using red, green, and blue excitation/emission channels, to visualize the CM-DiI-, DiD-, and DiO-labeled tracks and confirm probe placement in V1 and ACC.

Probe tracks were reconstructed using HERBS (Histological E-data Registration in rodent Brain Spaces)^38^. Images of each section were registered to the Allen Mouse Brain Common Coordinate Framework (CCFv3)^39^ at 10 µm resolution. The labeled track was then traced through the registered sections and fitted with a linear trajectory. Recording sites were assigned to brain regions from the position of the probe tip, so that each channel, and therefore each sorted unit, was assigned to an atlas-defined area.

### Video monitoring

Infrared lights and filters were placed in the box facing the mice. A camera (BFLY-U3-23S6M-C, FLIR) with zoom lenses (Zoom 700018–108mm, Navitar) acquired images synchronized with TTL pulses triggered by a FG085 miniDDSsquarefunction generator at 20 frames/s. The streams were saved as 8bit grayscale videos of 1980 × 1024 pixels. The right side of each animal, where large body, ear, paw, whisker, and tongue movements were visible, was recorded in a cropped frame of 1398 × 574 pixels. These recordings were analyzed using the DeepLabCut (DLC) behavioral tracking software.

## Data analysis

### Spike sorting and unit quality control

Electrophysiological recordings were automatically spike-sorted with Kilosort-2.5 (https://github.com/MouseLand/Kilosort)^40^. Sorted units were curated using automated quality-control criteria based on estimated noise-floor violations (the estimated proportion of spikes missed because they fell below the noise floor of the recording) and estimated refractory-period violations (the estimated proportion of spikes attributable to a different, non-primary neuron).

Quality-control accuracy was verified by manual review of a subset of units using the phy GUI (https://github.com/kwikteam/phy). For population decoding analyses, units were additionally required to pass probe-tracking confirmation of their location within V1 or ACC, and stability criteria applied to waveform amplitude, signal-to-noise ratio, and firing rate across the session; only units passing all criteria were retained.

### Data post-processing analyses

For peri-event time histograms (PETHs) aligned to task events (Figs. 2), neural activity was binned at 10 ms and smoothed with a causal half-Gaussian filter (SD = 50 ms). For the kernel-regression (GLM) analysis (Fig. 2), activity was binned at 10 ms, smoothed with a causal half-Gaussian filter (SD = 25 ms). PETHs shown for visualization include correct trials only; both correct and error trials were included in the GLM analysis. Units were included in the GLM analysis if their trial-averaged firing rate (in the window between stimulus and movement onset) was at least 0.1 spikes/s (Fig. 2D).

### Neural encoding model (GLM)

To characterize the neural encoding of task-relevant variables, we fitted a Generalized Linear Model (GLM) to the spiking activity of each neuron^41,42^. Spike trains were binned at 1 ms resolution using the neuroGLM toolbox (Pillow Lab, Princeton; github.com/pillowlab/neuroGLM). The design matrix was assembled from two categories of regressors—basis-expanded temporal filters and effects-coded categorical contrasts—as described below.

#### Trial selection

For each session, trials were drawn from both the core visual-attend and auditory-attend block periods with an additional 24-trial buffer appended to each block. Only trials containing either a 45° or 135° grating as the target or distractor stimulus were included (tr45 and tr135). Trials were ordered by orientation identity and block membership, yielding a set of trials spanning both block contexts and both grating orientations.

#### Design matrix

The design matrix X contained two structurally distinct types of regressors, assembled on a trial-by-trial basis at 1 ms resolution.

Basis-expanded temporal filters. Four task epochs were modelled with smooth temporal basis functions (raised cosine), which allowed the GLM to learn an arbitrary smooth time course of neural activity within each epoch rather than assuming a fixed boxcar shape: (1) pre-stimulus onset (0–200 ms from trial onset): 2 raised-cosine basis functions spanning 300 ms, capturing the pre-stimulus baseline and pre-stimulus ramp (start); (2) stimulus period (500–2500 ms): 8 raised-cosine basis functions spanning 2000 ms, capturing the full stimulus-driven response, with the basis covering the grating display window; (3) response/choice period (1950–3450 ms, i.e., 400 ms after choice onset to choice offset): 10 raised-cosine basis functions spanning 2000 ms, capturing motor and post-choice activity (response); and (4) outcome period (2500–4500 ms): 5 raised-cosine basis functions spanning 2000 ms, capturing reward- and punishment-related activity (outcome).

Effects-coded categorical contrasts. The remaining regressors each contributed a single column to the design matrix, encoding trial-level categorical variables as effects codes (contrast values) scaled by a fixed-duration boxcar within the relevant task epoch. These regressors were not z-scored, so that the fitted coefficient retained a direct interpretation as the change in log firing rate when that variable is active. All categorical regressors were encoded as contrast (+1/−1) or indicator (0/1) variables rather than pairs of complementary indicator variables, to ensure linear independence from the basis-expanded columns already spanning the same time windows.

### Model fitting

For each neuron, the binned spike train (1 ms bins) was smoothed with a 100-sample (100 ms) moving average and fit using a Poisson GLM with a log link function (MATLAB fitglm, ‘Distribution’,‘poisson’, ‘Link’,‘log’). Goodness of fit was assessed using the pseudo-R² (proportion of explained deviance):

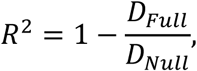

where *D_Full_* is the deviance of the full model and *D_Null_* is the deviance of an intercept-only model. Only neurons with R² > 0.05 were included in subsequent analyses.

### Covariate contribution analysis (LOCO)

The contribution of each covariate to neural encoding was quantified using a Leave-One-Covariate-Out (LOCO) approach. For each covariate k, a reduced model was fit by removing the corresponding column(s) from the design matrix and refitting. Two complementary statistics were computed from the full and reduced model pair:

1. Significance (selectivity): a likelihood-ratio test (LRT) was performed with test statistic^8^:

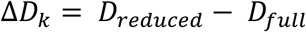

A neuron was classified as selective for covariate k at α = 0.05.

1. Relative contribution (proportional contribution): the proportional contribution of covariate k to a neuron’s total explained variance was computed as:

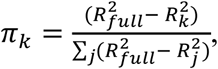

where R²_k_ is the pseudo-R² of the reduced model without covariate k and the sum in the denominator runs over all covariates j. This normalizes each covariate’s unique contribution relative to the total unique variance attributable to all covariates. Population-level proportional contribution was summarized as the mean ± SEM of π_k_ across neurons with R² > 0.05.

### Population decoding analyses

Neuropixels recordings from V1 and ACC spanned all cortical layers of V1 and were oriented orthogonal to the columnar axis of ACC. Spike-sorted units were selected based on probe-tracking coordinates and the stability criteria described above; all subsequent decoding analyses used units passing these criteria. For decoding, neural activity was binned at 1 ms resolution and smoothed with a 10 ms boxcar, then organized into trial-averaged firing-rate matrices; population responses were structured as three-dimensional arrays (trials × units × time bins) for each block within each session.

Each recording session consists of alternating VA and AV context blocks. Behavioral performance was tracked as a 10-trial moving average of rewarded trials, and expert-phase onset within a block was defined as the first trial at which this moving average reached or exceeded 0.85. All decoding analyses were restricted to expert-phase trials unless otherwise specified, so that neural representations were examined during periods of stable, context-appropriate behavior.

Linear support vector machine (SVM) decoders were trained to discriminate visual stimulus identity (target vs. distractor orientation) from population activity in sliding 40-trial blocks, advancing in steps of 5 trials. Each block decoder was trained and tested within the same trial block using 5-fold cross-validation (repeated twice), yielding a time-resolved decoding-accuracy matrix (trial blocks × time bins); signed decoder weight vectors were retained for all subsequent analyses. SVM was trained on 80% of trials and tested on 20% of trials for each iteration with no overlap of trained and tested sets. Mean decoding time courses were computed by averaging across expert-phase batches within each block and then across sessions (Fig. 3). Visual decoding accuracy was computed separately for VA and AV context blocks, and the VA−AV decoding-difference time course — the difference between mean VA and mean AV expert-batch accuracy at each time bin — was used as the primary measure of context-dependent modulation of visual stimulus coding (Fig. 3). All decoding used the MVPA-Light toolbox^43^ with a linear SVM kernel, k-fold cross-validation, and classification accuracy as the performance metric.

A single context decoder was trained to discriminate VA versus AV context from the same V1 and ACC populations. Expert-phase trials (≥85% task performance) from all available VA blocks and all available AV blocks within a session were pooled separately, balanced to equal size by random subsampling without replacement (fixed random seed, 42), and labelled VA = 1, AV = 2. Context decoding was performed with a linear SVM trained independently at each time bin, and the decoder was tested on trials excluded from its training set, with accuracy defined as the fraction of test trials classified as the correct context. The trial blocks from AV and VA session blocks were tested separately. This yielded a time-resolved context-accuracy time course for each test batch. Temporal generalization of the context representation was assessed with the corresponding time × time classification function using the same training and test sets, producing a [time bins × time bins] matrix in which entry (t_train_, t_test_) gives the accuracy of a decoder trained at t_train_ and tested at t_test_; broad off-diagonal structure indicates a stable, time-invariant context representation. Context decoder weights were extracted from the trained model parameters, yielding a signed weight matrix W_ctx_ [units × time bins]. Epoch-averaged context weights were computed as the mean absolute weight across an early stimulus epoch (0.5–2.5 s).

To characterize the progressive consolidation of context representations across trials within a session block, the pooled context decoder was applied to sliding 40-trial test blocks (step = 5 trials) within each VA and AV session block separately, with no trial block permitted to cross a session block boundary. Each trial block contained trials from a single context and was scored against that block’s true context label. To keep test trials out of the training set, expert-phase trials belonging to the current trial block were removed from the training pool, the pool was re-balanced between VA and AV by random subsampling, and the decoder was retrained before testing that trial block. For each trial block, the time-resolved context-decoding accuracy (train time = test time) was obtained. The early-epoch mean (0.5–2.5 s) accuracy per batch was extracted and plotted as a function of normalized trial position within the block (0 = block onset, 1 = block end).

The angle between the visual stimulus and context decoder hyperplanes (Fig. 4F) was computed at each time bin to characterize the geometric relationship between the two representations. At each time bin t, the composite visual weight vector w_vis_(t) (averaged across expert VA batches) and the context weight vector w_ctx_(t) were each normalized to unit length. The cosine similarity was defined as their dot product:

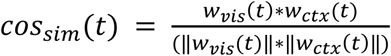

A value of 90° indicates orthogonal subspaces; values above or below 90° indicate anti-alignment or co-alignment, respectively. Grand-mean angle and cosine-similarity time courses were computed across sessions, and the mean angle during the early stimulus epoch was tested against 90° with a one-sample t-test across sessions.

### Statistical testing

Unless otherwise specified, all statistical comparisons used two-tailed tests. Multiple-comparison correction used the Bonferroni where multiple comparisons were made. Effects were considered significant at α = 0.05. Results are reported as mean ± SEM across sessions unless otherwise noted. All analyses were implemented in MATLAB (version R2025b) using custom code together with the neuroGLM^41^ and MVPA-Light^43^ toolboxes.

### Software

Spike sorting used Kilosort 2.5 (https://github.com/MouseLand/Kilosort) with manual curation in phy (https://github.com/kwikteam/phy). The GLM encoding-model pipeline was implemented in MATLAB using the neuroGLM toolbox (Pillow Lab; github.com/pillowlab/neuroGLM) for design-matrix construction and MATLAB’s fitglm for parameter estimation. Population decoding used the MVPA-Light toolbox. Computationally intensive analyses were run on the UCLA OARC Hoffman2 cluster.

## Acknowledgements

Daily training and other aspects of the behavioral experiments were carried out with the help of a dedicated team of talented UCLA undergraduate students. We thank Dr. João Couto for valuable suggestions regarding the experiments.

This work is dedicated to the memory of Helya Bahari-Kashani.

## Ethics declaration

The authors declare no competing interests.

## Data availability

Data and analysis code will be made available after manuscript acceptance. Any additional information to analyze the data reported in this paper will also be made available from the corresponding authors.

